# Analysis of mycobacterial infection-induced changes to host lipid metabolism in a zebrafish infection model reveals a conserved role for LDLR in infection susceptibility

**DOI:** 10.1101/250548

**Authors:** Matt D. Johansen, Elinor Hortle, Joshua A. Kasparian, Alejandro Romero, Beatriz Novoa, Antonio Figueras, Warwick J. Britton, Kumudika de Silva, Auriol C. Purdie, Stefan H. Oehlers

## Abstract

Changes to lipid metabolism are well-characterised consequences of human tuberculosis infection but their functional relevance are not clearly elucidated in these or other host-mycobacterial systems. The zebrafish-*Mycobacterium marinum* infection model is used extensively to model many aspects of human-*M. tuberculosis* pathogenesis but has not been widely used to study the role of infection-induced lipid metabolism. We find mammalian mycobacterial infection-induced alterations in host Low Density Lipoprotein metabolism are conserved in the zebrafish model of mycobacterial pathogenesis. Depletion of LDLR, a key lipid metabolism node, decreased *M. marinum* burden, and corrected infection-induced altered lipid metabolism resulting in decreased LDL and reduced the rate of macrophage transformation into foam cells. Our results demonstrate a conserved role for infection-induced alterations to host lipid metabolism, and specifically the LDL-LDLR axis, across host-mycobacterial species pairings.

**Funding:** This work was supported by the Australian National Health and Medical Research Council (APP1099912 and APP1053407 to S.H.O.); Meat and Livestock Australia (P.PSH. 0813 to A.C.P. and K. dS); the Marie Bashir Institute for Infectious Diseases and Biosecurity (grant to S.H.O., A.C.P. and K. dS); the Kenyon Family Foundation Inflammation Award (grant to S.H.O.); the University of Sydney (fellowship to S.H.O.); Consellería de Economía, Emprego e Industria (GAIN), Xunta de Galicia (grant IN607B 2016/12 to Institute of Marine Research (IIM-CSIC)).

## 1. Introduction

Mycobacterial manipulation of host lipid metabolism and the subsequent transformation of macrophages into foam cells are important pathways in the pathogenesis of *Mycobacterium tuberculosis* (*Mtb*) (Lovewell et al., 2016). Mycobacteria drive the transformation of infected macrophages into foam cells through the imbalance of low density lipoprotein (LDL) influx and efflux mechanisms (Cardona et al., 2009). Given the significant proportion of the mycobacterial genome that is allocated to lipid metabolism, and the utilisation of lipids such as cholesterol for growth *in vitro*, the abundance of neutral lipids contained within the granuloma microenvironment may supply a carbon source for intracellular mycobacterial survival and persistence (Pandey and Sassetti, 2008).

Dyslipidaemia and foam cell differentiation are known to contribute to disease pathology and prognosis in human *Mtb* infection where foamy macrophages may inhibit lymphocyte access to infected macrophages, increase the build-up of caseum at the center of granulomas, and provide lipids utilized during evasion of phagocytosis (Dong et al., 2017; Russell et al., 2009). However, the involvement of dyslipidemia in the pathogenesis of non-tubercular mycobacterial infections such as *Mycobacterium avium* subspecies *paratuberculosis* (MAP) or *Mycobacterium marinum*, and in non-rodent animals, has not been well explored. Our group has identified infection-induced changes to host lipid metabolism genes in MAP-infected cattle and aberrant host lipid metabolism gene expression and intracellular cholesterol accumulation within MAP-infected macrophages (Johansen et al., submitted; Thirunavukkarasu et al., 2014). Here, we have turned to the natural host-pathogen pairing of the zebrafish-*M. marinum* platform to investigate the conservation and consequences of altered host lipid metabolism as a conserved motif across mycobacterial infections.

Zebrafish (*Danio rerio*) have functional innate immune and digestive systems from early in embryogenesis with a high degree of homology to other vertebrates and are adaptable to the study of early host-pathogen interactions (Traver et al., 2003). *M. marinum* is a natural pathogen of fish and amphibian species and genomic analysis has identified *M. marinum* as the closest genetic relative of the *Mtb* complex, sharing 85% of orthologous regions with *Mtb* (Stinear et al., 2008). Zebrafish models of *M. marinum* infection have provided unique insight into host-pathogen interplay responsible for granuloma formation, exquisitely reproducing the cellular structure of the human-*M. tuberculosis* granulomas, including the presence of lipid-rich foam cells (Cronan et al., 2016; Johansen et al., 2018b; Oehlers et al., 2017; Oehlers et al., 2015).

The aim of this study was to explore the biological significance of infection-induced changes to lipid metabolism in a model non-tubercular mycobacterial infection. Our findings demonstrate the key lipid metabolism node Low Density Lipoprotein Receptor (LDLR) mediates the uptake of lipids into the mycobacterial granuloma niche and aids the transformation of macrophages into foam cells.

## 2. Materials and Methods

### 2.1. Zebrafish breeding and ethics statement

Adult zebrafish were housed at the Garvan Institute of Medical Research Biological Testing Facility (St Vincent’s Hospital AEC Approval 1511). Zebrafish embryos were obtained by natural spawning and embryos were raised at 28°C in E3 media. All experiments and procedures were completed in accordance with Sydney Local Health District animal ethics guidelines for zebrafish embryo research.

### 2.2. Morpholino design and microinjection

Splice-blocking morpholino oligonucleotide for *LDLR* (gene variant *ldlra*) was designed from Ensembl transcript sequences (http://www.ensembl.org/index.html) and purchased from Gene Tools. The *LDLR* morpholino (5’-ATCACATTTCATTTCTTACAGCAGT-3’) was targeted at the exon 4/intron 4 and 5 splice junction that was expected to cause the excision of exon 4. The control morpholino (5’-CCTCTTACCTCAGTTACAATTTATA-3’) is a negative control targeting a human beta-globin intron known to cause little phenotype variations in zebrafish.

Embryos at the 1–4 cell stage were injected with 2 nL of morpholino into the yolk. Embryos were raised at 28°C in E3 media.

### 2.3. Molecular analysis of *ldlra* knockdown

RNA was extracted in TRIZOL (Thermofisher) by precipitation and cDNA was synthesised with an Applied Biosystems High Capacity cDNA reverse transcription kit (Thermofisher). PCR was performed with *ldlra*-specific primers (5’-AGAGCTGGAAATGTGACGGA-3’ and 3’-CTCATCTGGACGGCATGTTG-5’) and visualised by gel electrophoresis.

### 2.4. Oil Red O staining

Oil Red O lipid staining on whole mount embryos was completed as previously described (Johansen et al., 2018b; Passeri et al., 2009). Briefly, embryos were individually imaged for bacterial distribution by fluorescent microscopy, fixed, and stained in Oil Red O (0.5% w/v in propylene glycol). Oil Red O staining intensity at sites of infection were quantified in ImageJ and calculated as the pixel density above background tissue.

### 2.5. Infection of zebrafish embryos

At 30 hours post-fertilisation, zebrafish embryos were anaesthetised with 160 µg/mL tricaine (Sigma-Aldrich) and infected with approximately 200 CFU fluorescent M strain *M. marinum* via caudal vein injection.

### 2.6. Lipoprotein assay

Low density lipoprotein abundance was measured using the HDL & LDL/VLDL Cholesterol Assay Kit (Cell Biolabs) as per manufacturer’s instructions and based on previously published methods (O’Hare et al., 2014). Briefly, 10 embryos per treatment group were terminally anaesthetised and placed into BHT solution (10 µg/mL, Sigma-Aldrich) and homogenised. Homogenates were centrifuged at 4°C at 2000 x *g* for 5 mins to pellet debris. The supernatant was mixed with an equal volume of precipitation solution to separate the high density lipoprotein (HDL) and low density lipoprotein (LDL) fractions. Each lipoprotein fraction was mixed with an equal volume of reaction mix and quantified by fluorescent plate reader.

A standard curve was included in each run to calculate the concentration of lipoproteins within each treatment, however the measured quantities of LDL varied considerably between between replicates (eg control experiment 1 vs control experiment 2) so we present data as arbitrary normalised units to facilitate fold change comparisons between experiments.

### 2.7. Imaging

Live zebrafish embryos were anaesthetized in M-222 (Tricaine) and mounted in 3% methylcellulose for imaging on a Leica M205FA fluorescence stereomicroscope. Further image manipulation and/or bacterial quantification was carried out with Image J Software Version 1.51j.

### 2.8. Quantification of *M. marinum* burden by fluorescent pixel count

Infection burden was measured as the number of pixels in each embryo above background fluorescence in ImageJ (National Institutes of Health) and pixels counted using the ‘Analyse particles’ function (Matty et al., 2016).

### 2.9. Quantitative PCR

Quantitative PCR (qPCR) reactions were completed as previously described (Taylor et al., 2008) using Mx3000P real-time PCR systems (Stratagene). Gene-specific primers were generated for *abca1* (5’-AGCTGCTGGTCGAGATCATA-3’ and 3’-CTGTCTCACTGCATGCTTGG-5’) for comparison against housekeeping gene *actb* (5’-CCTTCCAGCAGATGTGGATT-3’ and 3’-CACCTTCACCGTTCCAGTTT-5’). Samples were run in duplicate with reaction specificity confirmed based on the post-amplification dissociation curve.

### 2.10. Statistical analysis

Graphpad Prism was used to perform statistical testing as indicated.

## 3. Results

### 3.1. Infection-induced altered host dyslipidemia is recapitulated in the zebrafish-*M. marinum* infection model

Previous reports have identified hypercholesterolemia as a correlate of disease pathology during human-*Mtb* and ruminant-*M. avium* paratuberculosis infections (Dong et al., 2017). To determine whether these findings were conserved within the zebrafish-*M. marinum* platform, we infected zebrafish embryos with *M. marinum* then analysed the total embryo LDL content at 5 dpi. Calculation of lipoprotein content revealed an increase in LDL demonstrating conservation of the infection-induced dyslipidemia phenotype in our zebrafish embryo model (Figure 1A).

**Figure 1:**
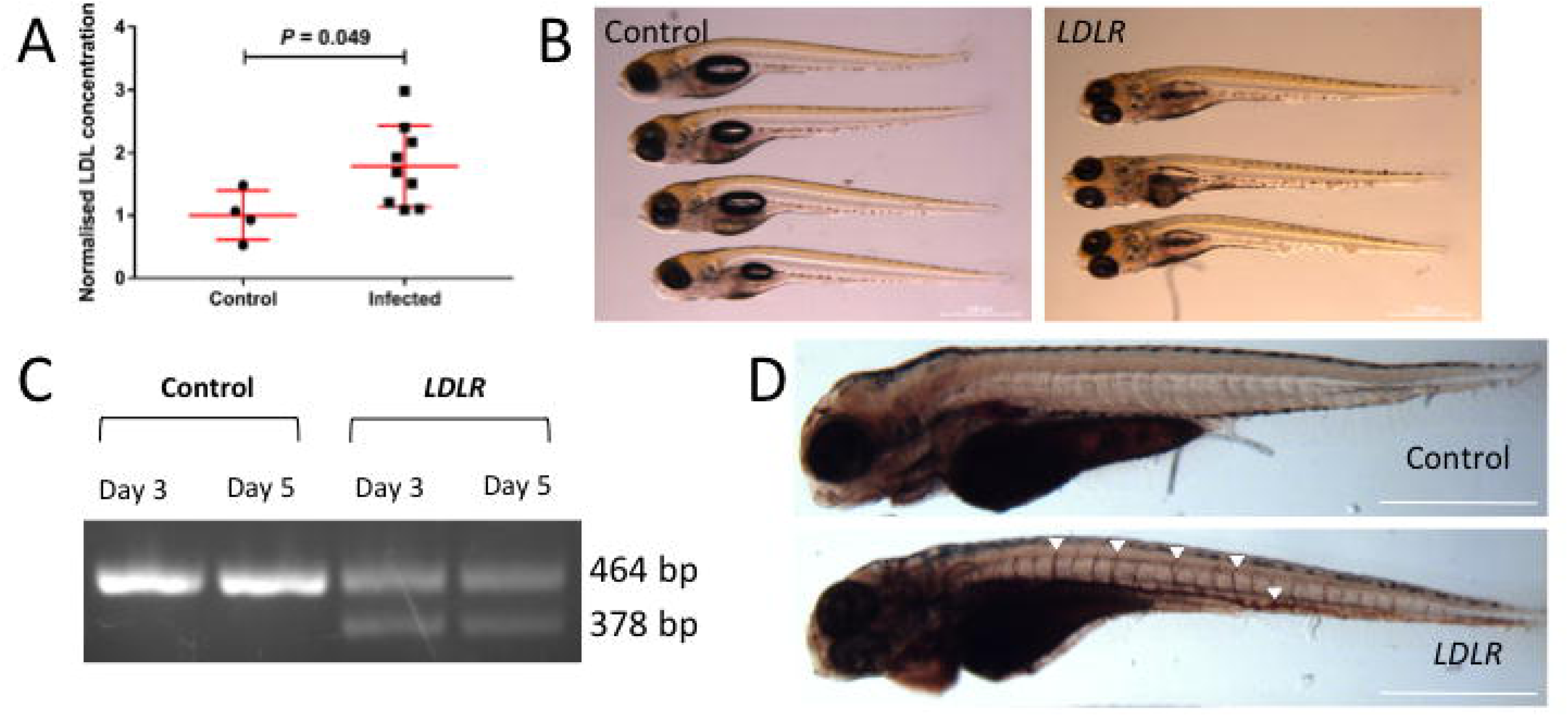
LDL is increased by infection and validation of Ldlra knockdown. *A*, Quantification of LDL concentration in wild-type zebrafish embryos at 5 days following *M. marinum* infection. Each data point represents a pooled sample of 10 embryos. Scale bars represent 500 µm. Statistical tests were performed using unpaired Student’s t-tests. Experiments were completed with 4 independent biological replicates. *B*, Normal morphology and phenotypes at 5 days post-fertilisation as visible by light microscopy. *C*, Molecular validation of *ldlra* morpholino efficacy at 3 and 5 days post-treatment demonstrating depletion of native 464 bp amplicon and appearance of aberrant 378 bp amplicon. *D*, Oil Red O staining demonstrates visible lipid abundance within the vasculature of Ldlra-deficient embryos. White arrows represent expansive lipid accumulation within the vasculature.

### 3.2. Knockdown of *Ldlra* reduces *M. marinum* burden in zebrafish embryos

Infection of macrophages with various species of mycobacteria, including in the zebrafish-*M. marinum* system, increases *LDLR* transcription (Cronan et al., 2016; Johansen et al., submitted; Kim et al., 2010). Take together with our observation of infection-induced changes in the ligand LDL in infected zebrafish embryos, we tested the functional relevance of LDLR by morpholino knockdown of the zebrafish *ldlra* gene (O’Hare et al., 2014). As previously reported by O’Hare et al., knockdown of Ldlra had no effect on embryo development and editing of the *ldlra* transcript was observed (Figures 1B and 1C). Oil Red O staining for lipids in Ldlra-depleted embryos revealed marked vascular lipid accumulation consistent with previous reports of Ldlra depletion being atherogenic (Figure 1D) (O’Hare et al., 2014).

Following Ldlra-depletion, embryos were infected with fluorescent *M. marinum* and bacterial burden was analysed by fluorometry. Zebrafish embryos depleted of Ldlra displayed a significantly decreased bacterial burden at 3 and 5 dpi when compared to control embryos (Figure 2).

**Figure 2:**
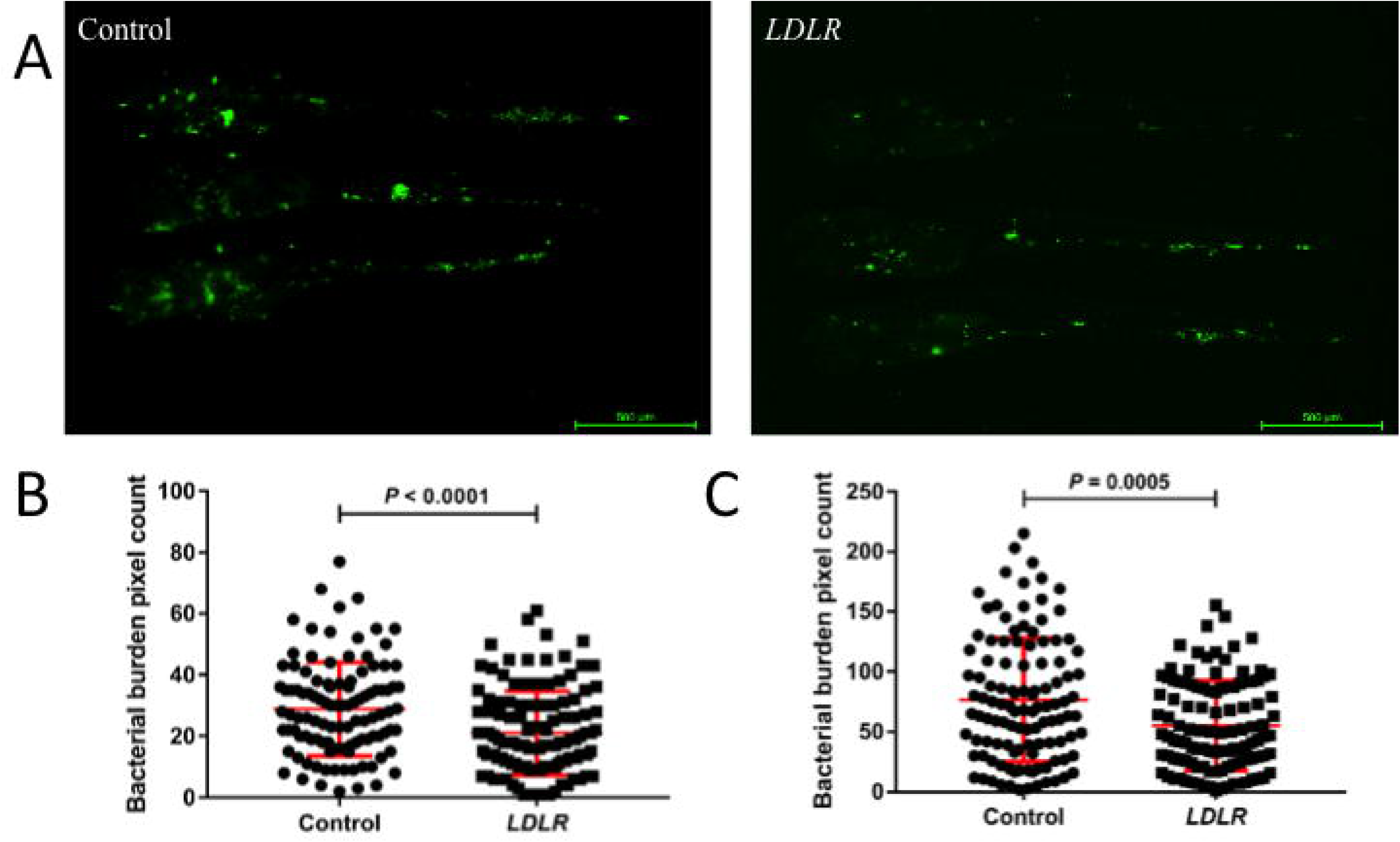
Knockdown of Ldlra reduces *M. marinum* bacterial burden. *A*, Representative images of *M. marinum* bacterial burden at 5 days-post infection. Scale bars represent 500 µm. *B*, Quantification of *M. marinum* bacterial burden at 3 days post infection. *C*, Quantification of *M. marinum* burden at 5 days post infection. Error bars represent standard deviations, each data point represents a single embryo. Analysis of *M. marinum* bacterial burden was unmatched T tests. Experiments were completed with 4 independent biological replicates for each panel.

### 3.3. Knockdown of Ldlra corrects infection-induced altered host lipid metabolism

Host lipid metabolism is a tightly controlled biological pathway, responsible for addressing the cellular lipid requirements through the regulation of influx and efflux mechanisms. Of the many surface receptors involved in lipid metabolism, LDLR is arguably the most important, playing a key role in driving the uptake of circulating LDL (Cardona et al., 2009). We hypothesised knockdown of Ldlra would thus restore homeostatic lipid metabolism.

Following *M. marinum* infection, expression of the LDLR-network and lipid efflux gene *ABCA1* was markedly increased (Figure 3A). Depletion of Ldlra significantly reduced the magnitude of infection-induced *abca1* transcription (Figure 3A).

**Figure 3:**
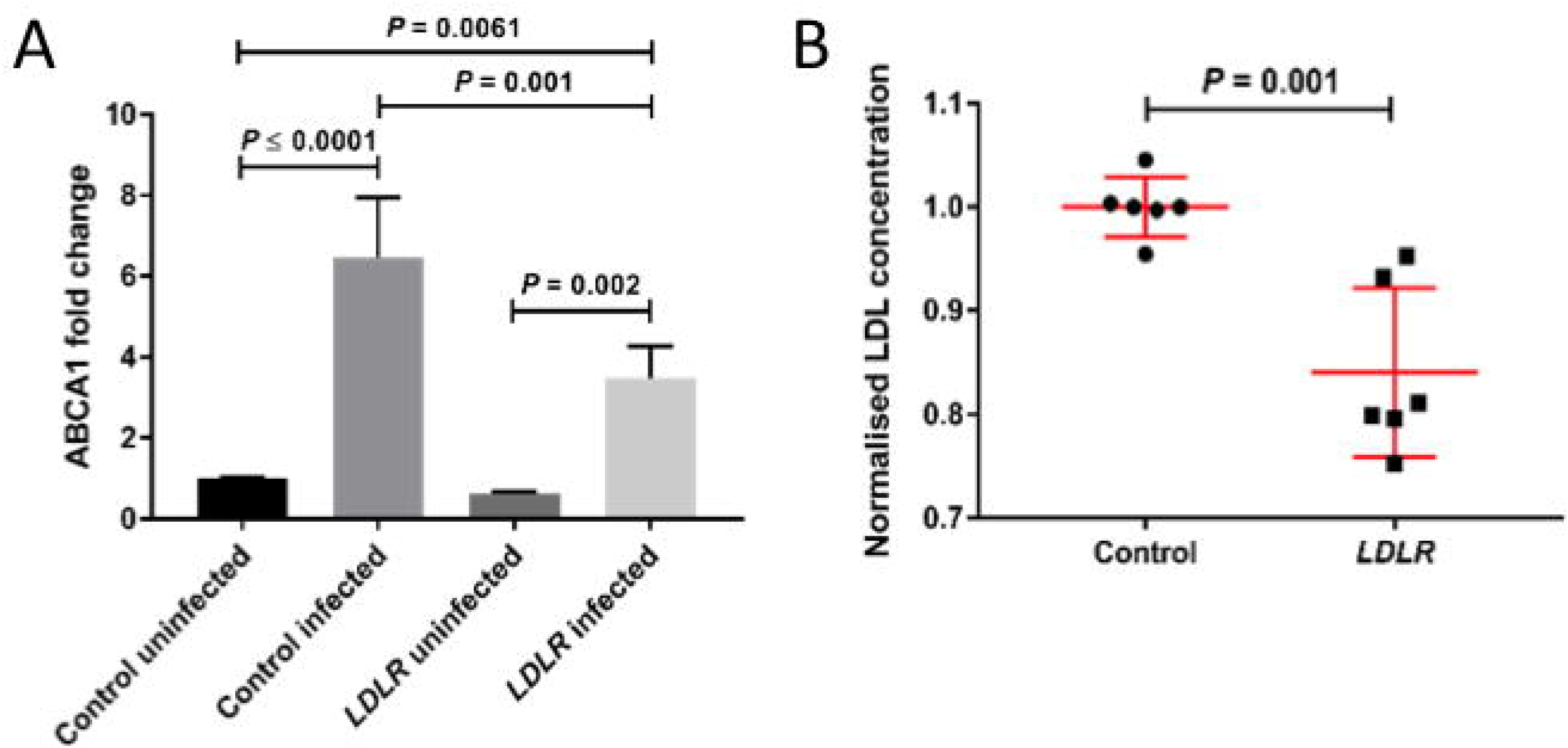
Mycobacterial infection-induced modulation of host lipid metabolism is corrected in Ldlra-deficient embryos. *A*, *abca1* transcript fold change expression at 5 days post-infection measured by qPCR, normalised to control uninfected sample. Each column represents two independent biological replicates analysed with technical replicates. *B*, Quantification of LDL concentration in Ldlra-deficient embryos at 5 days-post infection. Each data point represents a pooled sample of 10 embryos completed with four independent replicates. Gene expression analysis was completed using one-way ANOVA. Analysis of LDL concentration was completed using matched Student’s *t*-tests.

Next, we hypothesised that depletion of LDLR would prevent infection-induced hypercholesterolemia. Following *M. marinum* infection, Ldlra-deficient embryos displayed a decreased LDL compared to infected controls at 5 dpi (Figure 3B). Together, these data demonstrate conservation and correction of an infection-induced lipid metabolism network in our model.

At the cellular-to-tissue level, we hypothesised that the combination of reduced potential for LDL uptake and correction of total cholesterol levels would reduce granuloma lipid accumulation within granuloma foam cells in Ldlra-depleted embryos. Granuloma Oil Red O lipid staining density was reduced in infected Ldlra-deficient embryos compared to infected controls demonstrating a functional role for LDLR in directing bacterial-beneficial lipid accumulation during mycobacterial infection (Figure 4).

**Figure 4:**
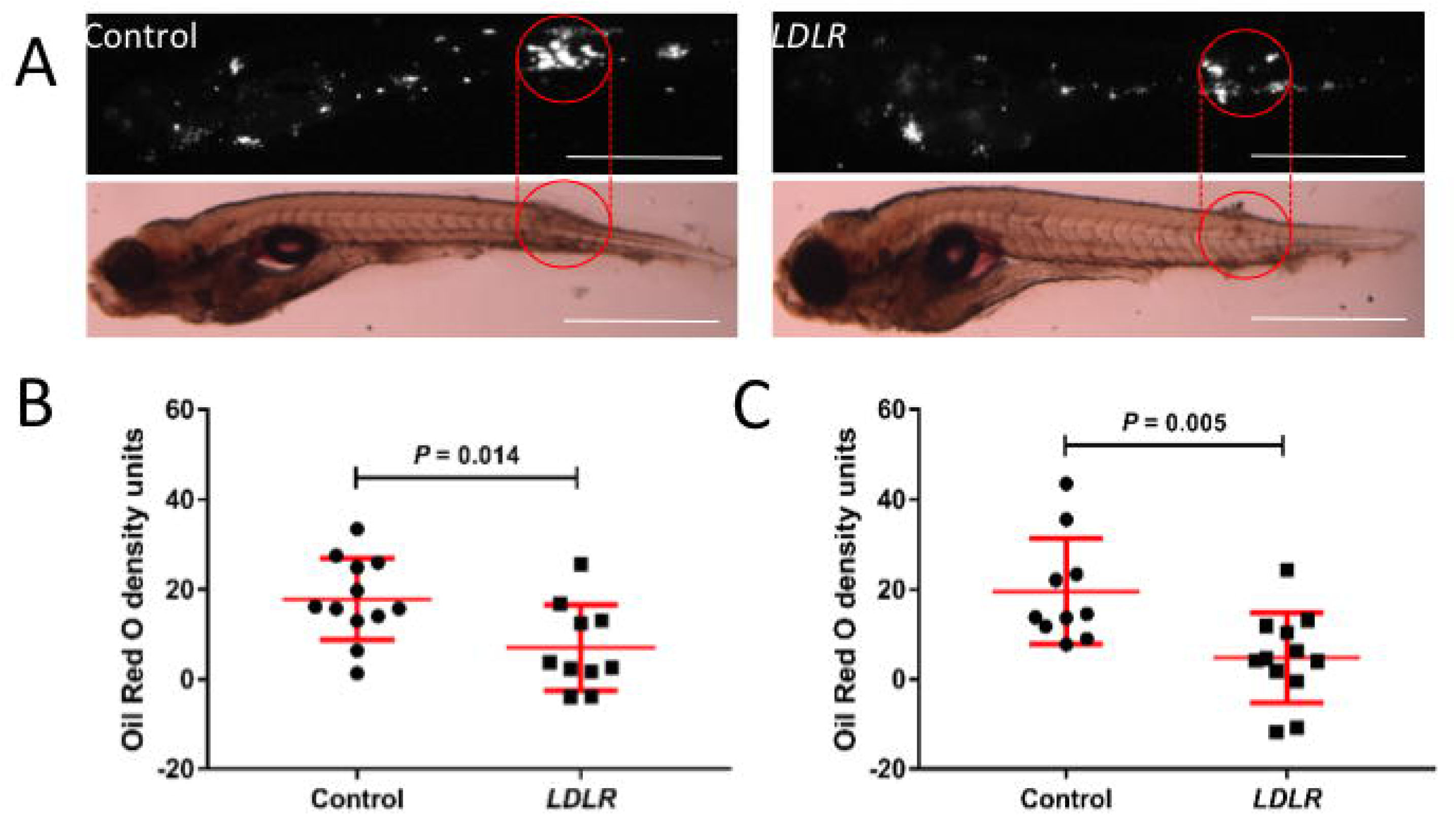
Infection-induced granuloma lipid accumulation is reduced in Ldlra-deficient embryos. *A*, Representative images of Oil Red O staining density at 5 days post-infection. Red circles indicate areas of embryos with largest foci of infection. Scale bar represents 500 µm. *B*, Quantification of Oil Red O density units at 3 days post-infection with *M. marinum*. *C*, Quantification of granuloma Oil Red O staining density units at 5 days post-infection. Each data point represents an individual granuloma. Error bars represent standard deviation. Statistical tests were performed using unpaired Student’s t-tests.

## 4. Discussion

The elevation of host lipids following mycobacterial infection has been well documented in both human and murine *Mtb* infections (Dong et al., 2017; Martens et al., 2008). Recently, we have shown that elevated serum cholesterol is associated with histopathological lesions in sheep and cattle infected with MAP (Johansen et al., 2018a). Here we show results in agreement with previous findings and suggest that the induction of host hypercholesterolemia is a conserved motif of pathogenic mycobacteria outside of mammalian hosts. Our findings in this study demonstrate 1) infection-induced altered host lipid metabolism is a conserved feature of non-tuberculous mycobacterial infection, and 2) host expression of LDLR is co-opted by mycobacterial infection to increase susceptibility to mycobacterial infection. Furthermore, our results demonstrate that *LDLR* knockdown significantly alters the availability of lipoproteins and decreases granuloma Oil Red O staining density, providing strong evidence to suggest that the abundance of neutral lipids and therefore foam cells, was largely decreased as a consequence of *LDLR* depletion.

Host lipid metabolism is a complex pathway involving many transcription factors, surface molecules and feedback mechanisms. In the absence of key components, compensatory pathways are utilised to achieve cellular homeostasis. Expression of *abca1* was increased following *M. marinum* infection, in agreement with previous studies identifying the upregulation of *ABCA1* by mycobacterial infection across multiple host species (Long et al., 2016). Interestingly, there was a notably lower expression in *ldlra*-deficient embryos, presumably due to the decreased efflux of lipids from mycobacteria-infected macrophages as a consequence of the altered lipoprotein abundance.

An important limitation of our study is the use of a single morpholino targeting the *ldlra* transcript in the embryo stage of development. Additional work with mutant adult zebrafish is necessary to fully elucidate the role of the LDL-LDLR axis in mycobacterial infection in the context of the adaptive immune system (Liu et al., 2018). The adult zebrafish platform will also allow finer control of available lipid pools through regulation of diet.

Collectively, this study provides evidence of conserved infection-induced altered host lipid metabolism in the zebrafish-*M. marinum* model, and highlights a crucial role of *LDLR* in infection susceptibility across host and pathogen species.

## Author contributions

M.D.J. Planned experiments, performed experiments, analysed data, wrote the paper; E.H. Planned experiments, performed experiments; J.A.K. Performed experiments; A.R. Performed experiments, contributed reagents; B.N. Performed experiments, contributed reagents; A.F. Contributed reagents; W.J.B. Analysed data, contributed reagents; K.dS. Planned experiments, analysed data, contributed reagents; A.C.P. Planned experiments, analysed data, contributed reagents; S.H.O. Planned experiments, performed experiments, analysed data, contributed reagents, wrote the paper.

## Acknowledgements

Drs Anneliese Ashurst and Gayathri Nagalingam for training assistance with laboratory equipment; Dr Kristina Jahn and Sydney Cytometry for assistance with imaging equipment. Special thanks goes to the animal house staff at the Garvan Institute of Medical Research and CSIC.

